# Comparison of vibrotactile and joint-torque feedback in a myoelectric upper-limb prosthesis

**DOI:** 10.1101/623926

**Authors:** Neha Thomas, Garrett Ung, Colette McGarvey, Jeremy D. Brown

## Abstract

**Background:** Despite the technological advancements in myoelectric prostheses, body-powered prostheses remain a popular choice for amputees, in part due to the natural sensory advantage they provide. Research on haptic feedback in myoelectric prostheses has delivered mixed results. Furthermore, there is limited research comparing various haptic feedback modalities in myoelectric prostheses. In this paper, we present a comparison of the feedback intrinsically present in body-powered prostheses (joint-torque feedback) to a commonly proposed feedback modality for myoelectric prostheses (vibrotactile feedback). In so doing, we seek to understand whether the advantages of kinesthetic feedback present in body-powered prostheses translate to myoelectric prostheses, and whether there are differences between kinesthetic and cutaneous feedback in prosthetic applications.

**Methods:** We developed an experimental testbed that features a cable-driven, voluntary-closing 1-DoF prosthesis, a capstan-driven elbow exoskeleton, and a vibrotactile actuation unit. The system can present grip force to users as either a flexion moment about the elbow or vibration on the wrist. To provide an equal comparison of joint-torque and vibrotactile feedback, a stimulus intensity matching scheme was utilized. Non-amputee participants (n=12) were asked to discriminate objects of varying stiffness with the prosthesis in three conditions: no haptic feedback, vibrotactile feedback, and joint-torque feedback.

**Results:** Results indicate that haptic feedback increased discrimination accuracy over no haptic feedback, but the difference between joint-torque feedback and vibrotactile feedback was not significant. In addition, our results highlight nuanced differences in performance depending on the objects’ stiffness, and suggest that participants likely pay less attention to incidental cues with the addition of haptic feedback.

**Conclusion:** Even when haptic feedback is not modality matched to the task, such as in the case of vibrotactile feedback, performance with a myoelectric prosthesis can improve significantly. This implies it is possible to achieve the same benefits with vibrotactile feedback, which is cheaper and easier to implement than other forms of feedback.

## 1 Background

Although upper-limb myoelectric prosthesis technology has advanced rapidly in recent years, the functionality of commercially available prostheses is still lacking compared to the natural limb. Among the listed features that amputees still desire in their myoelectric prosthesis are improved gripping, slip control, and sensory feedback [1]. For current clinically available myoelectric prostheses, sensory feedback from the terminal device is, with little exception, nonexistent. Amputees must therefore rely heavily on vision to complete activities of daily living, resulting in cognitive fatigue, as well as muscle fatigue due to overcompensation of grip force in object manipulation tasks [1, 2]. Endowing myoelectric prostheses with haptic feedback would help mitigate these issues as well as promote embodiment of the device, thereby allowing amputees to accurately and smoothly handle tasks like grasping, moving, and manipulating objects with fine dexterous control.

Despite the technological and aesthetic advantages available in myoelectric prostheses, body-powered prostheses remain a common choice for amputees [3]. This is due in part to the inherent kinesthetic (force and motion based) feedback provided by their mechanical linkages and cabling [4], which supports the concept of extended physiological proprioception [5, 6]. Specifically, grip force of the terminal device is displayed to the amputee as a torque about the shoulder joint. As empirically validated previously by members of our research group, the added utility of the kinesthetic (joint-torque) feedback available in body-powered prostheses provides a natural sensory advantage over traditional myoelectric prostheses [7].

For myoelectric prostheses, extensive research has been conducted on the utility of haptic devices as a sensory substitute for intact haptic sensation. One of the most commonly proposed modes of haptic feedback for upper-limb myoelectric prostheses is vibrotactile stimulation, whereby the grip force or aperture of the terminal device is mapped to the amplitude and/or frequency of the vibration stimulus [8]. Several papers have concluded that vibrotactile feedback is useful to enhancing performance in a myoelectric prosthesis. Witteveen *et al* showed that the addition of vibrotactile feedback increased stiffness discrimination accuracy 35% over chance [9]. More recently, Raveh *et al* found that vibrotactile feedback decreased the time to complete object manipulation tasks compared to no feedback in a myoelectric prosthesis [10]. Vibrotactile feedback has also been shown to be useful when operating a myoelectric prosthesis with limited or disturbed vision [11]. In a modified box and blocks task, contact cues displayed with vibration significantly reduced the number of broken blocks handled by transradial amputees [12]. In addition to vibrotactile feedback, other cutaneous haptic modalities such as electrotactile, pressure, and skin stretch have been shown to aid object manipulation performance with a myoelectric prosthesis [13–15].

Despite these findings, some investigations into the utility of haptic feedback have shown no benefit or improved functionality. In previous research from members of this research group, Brown *et al* found no significant improvement with either joint-torque feedback or vibrotactile feedback over no feedback in a grasp-and-lift task with a myoelectric prosthesis [16]. Similarly, no significant improvement in myoelectric control was found for a compensatory tracking task with pressure feedback over visual feedback [17]. Markovic *et al* showed that there was no significant advantage with a vibrotactile bracelet in a box and blocks task [18]. Vibrotactile feedback was also shown not to have a significant effect on grip force accuracy at medium and high grip forces [19]. Given the mixed results on the benefits of haptic feedback for myoelectric prostheses, it becomes clear why haptic feedback of any modality has remained largely absent from commercial prostheses.

Perhaps one of the factors contributing to the ambiguity regarding the utility of haptic feedback is the lack of proper comparisons between various haptic modalities. While comparisons can be made between different studies, the experimental investigations often involve non-standardized experimental hardware, different subject populations, or differing experimental tasks [13, 20, 21]. Even if the modality of feedback is the same, the manner in which it is presented can differ significantly [8, 18]. Together, these factors make any such comparison quite complicated. Even when various haptic modalities are compared within a single research study, such a comparison carries with it the confound that differences in performance can be attributed to differences in the perceived intensity of a stimulus rather than its somatosensory encoding. Additionally, as the research on haptic feedback for myoelectric prostheses heavily leans towards cutaneous feedback, it is unclear how the kinesthetic (joint-torque) feedback inherent to body-powered prostheses compares to the tactile feedback methods often proposed for myoelectric prostheses.

In this manuscript, we present a generalized methodology for comparing various haptic feedback modalities in a myoelectric prosthesis for the same experimental task. In our initial investigation, we compare joint-torque feedback to vibrotactile feedback. While joint-torque feedback may have practical limitations regarding the performance of activities of daily living, it is still useful for drawing comparisons between body-powered and myoelectric prostheses. We begin with an overview of our prosthesis testbed, followed by a discussion of the stimulus intensity matching scheme we employed to place both modalities on equal footing in terms of perceived stimulus intensity. Next, we describe and present the results of an empirical investigation in which we asked participants to discriminate between objects of varying stiffness in three separate conditions: no haptic feedback, joint-torque feedback, and vibrotactile feedback. In this study, we hypothesized that both haptic feedback conditions would outperform no haptic feedback, in line with the previously mentioned literature. In addition, since stiffness discrimination is a force-based task, we further hypothesized that joint-torque feedback would result in greater discrimination accuracy than vibrotactile feedback because of the modality similarities between the force measured by our prosthesis and the torque displayed to the user. Finally, we discuss the results of our study and comment on their broader implications for myoelectric prosthesis development.

## 2 Methods

### 2.1 Participants

We investigated the ability of n=12 able-bodied individuals (7 male, 5 female, age = 23.7*±*4.9 years) to discriminate objects in a 2-AFC task using a custom myoelectric prosthesis in three different feedback conditions. The duration of the experiment was approximately 90 minutes and participants were compensated at a rate of $10 per hour. All participants were consented according to a protocol approved by the Johns Hopkins School of Medicine Institutional Review Board (Study# IRB00147458). Participants were pseudo-randomized into two groups in an alternating fashion.

### 2.2 Experimental Apparatus

Our experimental prosthesis testbed consists of a custom 1-DoF voluntary-closing prosthetic terminal device mated to a custom prosthetic socket via a Hosmer Quick Disconnect Wrist as shown in Fig. 1a. The custom socket is designed to be worn by non-amputee participants. The prosthetic terminal device was driven by a custom linear actuator drive (see Fig. 1b). The actuator drive consists of a linear ballscrew (2mm lead) coupled to a Maxon RE50 (200 W) motor. The motor features a US Digital optical encoder (1024 CPR) and was driven by a Maxon ESCON 70/10 servocontroller in current-control mode with a gain of (1A/V). A Transducer Techniques LSP-10 10 Kg loadcell was attached to the ballnut of the ballscrew through a custom 3D printed carriage. The carriage was attached to a linear slide which allowed it to move freely with the ballnut. This linear drive provided pulling actuation on a Bowden cable, which in turn closed the terminal device.

**Figure 1.**
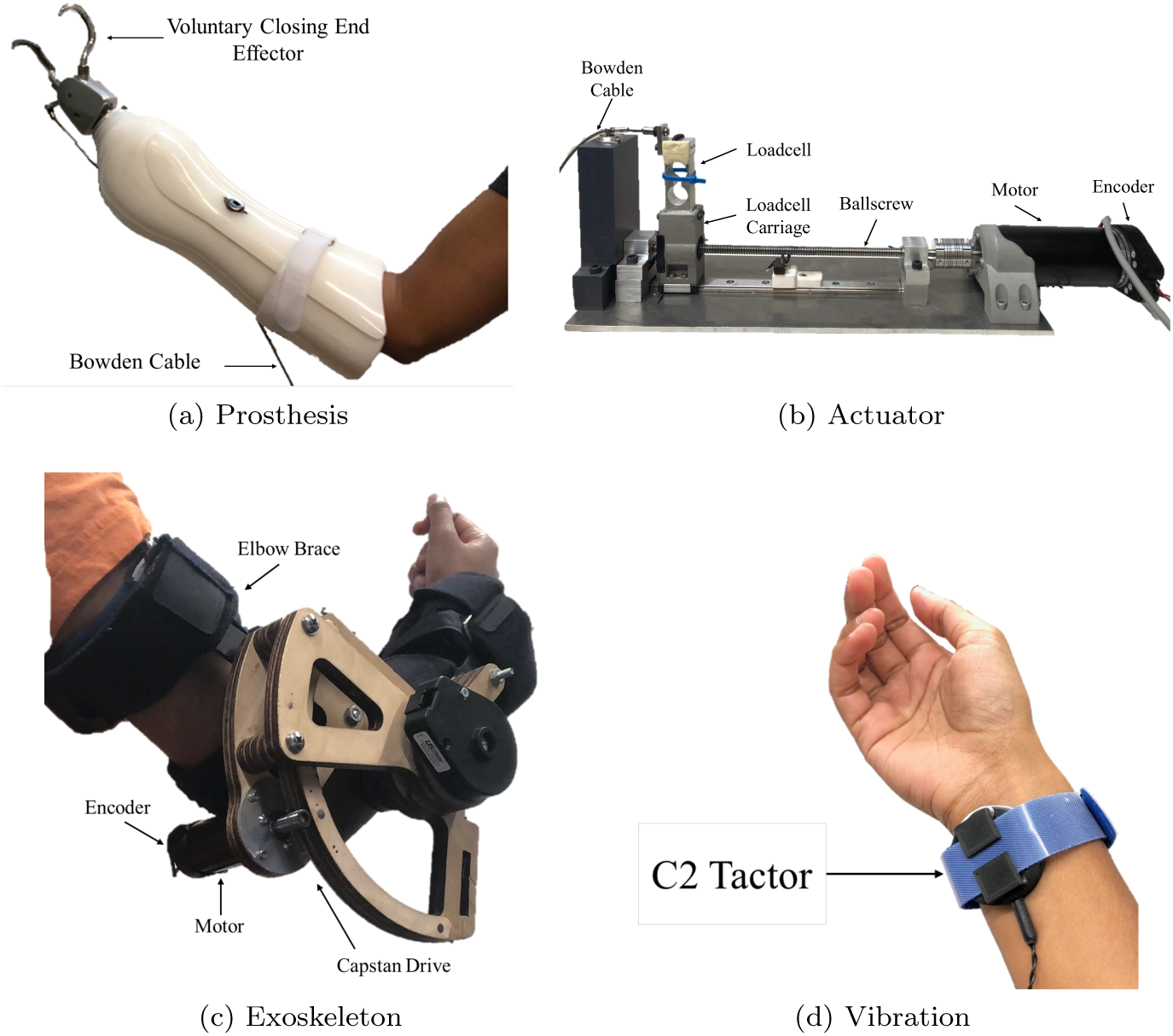
(a) Prosthesis terminal mated to custom socket for able-bodied individuals. (b) Linear actuator drive features a motor, encoder, ballscrew, and loadcell. The linear actuator drive connects to the prosthesis via a Bowden cable. (c) Capstan-driven exoskeleton used for applying joint-torque feedback, (d) C2-Tactor for applying vibrotactile feedback in a 3D printed housing with Velcro strap for wrist attachment

Joint-torque feedback was provided through a powered exoskeleton consisting of an Aircast Mayo Clinic elbow brace and a capstan drive featuring a Maxon RE30 (60 W) DC motor as shown in Fig. 1c. The motor features a Maxon encoder (1024 CPR) and was driven by a Maxon ESCON 70/10 servocontroller in current-control mode with a gain of (1A/V). The angular position of the elbow joint was measured with a US Digital optical encoder (1024 CPR). The exoskeleton was capable of delivering a maximum torque of 3.24 Nm as a flexion moment at the elbow joint. This exoskeleton was originally presented in previous work by members of our research group, Brown *et al* [7]. An explanation of the joint-torque stimulus is detailed in Section 2.6.4.

Vibrotactile feedback was provided by a C2-Tactor powered by a programmable Tactor Control Unit (Engineering Acoustics, Inc.) The C-2 tactor was housed in a 3D printed enclosure and secured to the participant’s wrist with a Velcro wristband (see Fig. 1d). The C-2 Tactor is capable of producing a maximum skin displacement of 0.499 mm. An explanation of the vibration stimulus is detailed in Section 2.6.4.

Electromyographic (EMG) signals were acquired from the left wrist flexor and extensor muscle group using a Delsys Bagnoli 16-channel EMG system with two surface electrodes.

Data acquisition and control were implemented through a Quanser QPIDe DAQ operating at a 1 kHz sampling rate. The entire system was controlled by a Dell Precision T5810 desktop running MATLAB R2017a with the Simulink Desktop Real-Time environment along with Quanser’s QUARC real-time Simulink blockset and a custom block containing the Engineering Acoustics API for C-2 tactor control.

### 2.3 Cross Modal Matching

Prior to performing the experimental task, we performed a cross-modal matching task based on the paradigm developed by Pitts *et al* to calibrate the two haptic modalities, vibrotactile and joint-torque feedback, in terms of their stimulus intensity [22]. This step is done so that any difference in experimental performance between the two modalities is the result of differences in the sensory encoding and not differences in the perceived stimulus intensity. The participant was instructed to sit on a stool facing a monitor. The participant then donned the exoskeleton and tactor wristband on their right arm with the assistance of the experimenter, and was then instructed to rest their right arm on an armrest.

#### 2.3.1 Phase 1 – Intensity Exploration

Participants were presented with a graphical user interface (GUI) (Fig. 2a) with adjustable sliders that allowed them to explore the range of vibrotactile and joint-torque stimulus intensities. For joint-torque feedback, participants were told that they could either passively let the exoskeleton flex their elbow, or move their arm back and forth to feel the torque generated by the exoskeleton. They were asked to use the same method during the experimental task; the method used was recorded in post-condition surveys. During this intensity exploration phase, both vibration and joint-torque cues played simultaneously for a period of four seconds. The entire phase lasted for a maximum duration of two minutes.

**Figure 2.**
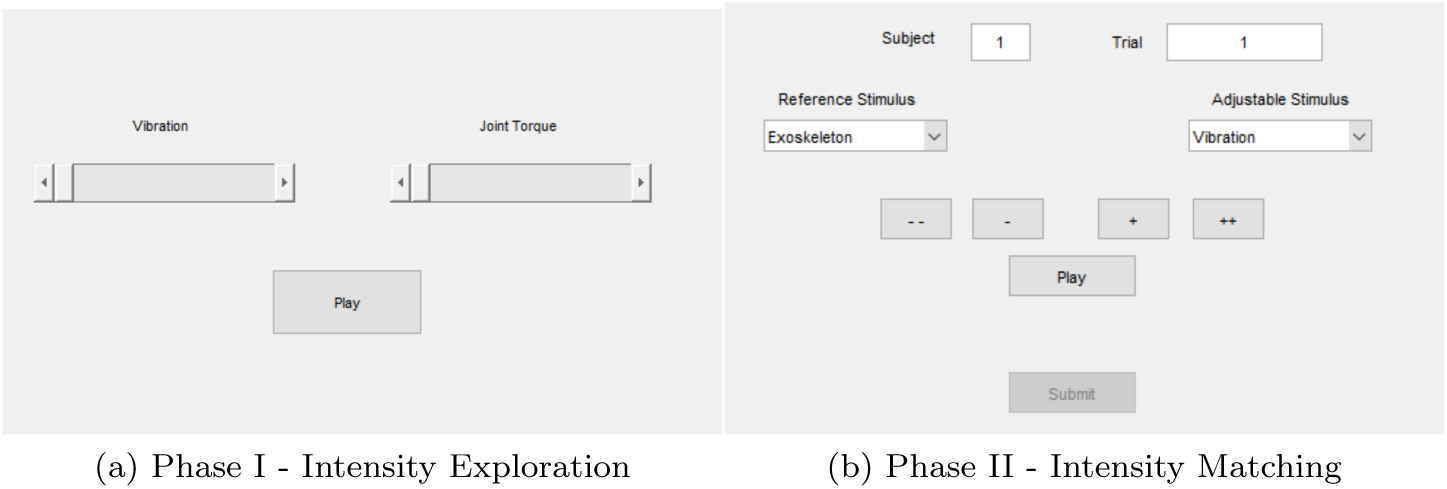
GUIs for the (a) intensity exploration and (b) intensity matching phases in the cross-modal matching step.

#### 2.3.2 Phase 2 – Intensity Matching

After participants were comfortable with the range of stimulus intensities, they were presented with a second GUI (Fig 2b). Participants were then given a reference torque stimulation and were instructed to adjust the vibration stimulation using the GUI pushbuttons until they perceived the two intensities to be the same. Participants matched four distinct reference joint-torque values three times each, for a total of twelve matches. They were given a maximum of two minutes per match. If two minutes was exceeded, the participant submitted their current match value. The four reference values were 0.66, 1.43, 2.08, and 2.93 Nm, which correspond to values of 0.208, 0.433, 0.631, and 0.892 when normalized with respect to the exoskeleton’s maximum torque. These four values were chosen to acquire data at the extreme ends as well as get points on either side of the halfway point. Through preliminary experimental investigation, it was decided that four reference values would be sufficient to increase the curve resolution without significantly lengthening the total experiment duration.

After completing the matching, the average match values were plotted against the four reference points. A fifth point was used to indicate a minimum reference joint-torque stimulus matched with the minimum easily perceived vibrotactile stimulus. These thresholds were determined through four pilot experiments with the method of limits and are are all at least 15 dB above the true absolute thresholds reported in [23] and [24]. This first point was the same value for all participants. A two-parameter exponential function was fit to the data to approximate the participant’s cross-modal curve. This function was determined to work best through a heuristic approach. One such example graph is shown in Fig. 3. Other participants exhibited similar trends but with different values. In the experiment, this function was used to convert exoskeleton motor commands to the corresponding vibration commands during the vibrotactile feedback condition as described in Section 2.6.4 below.

**Figure 3.**
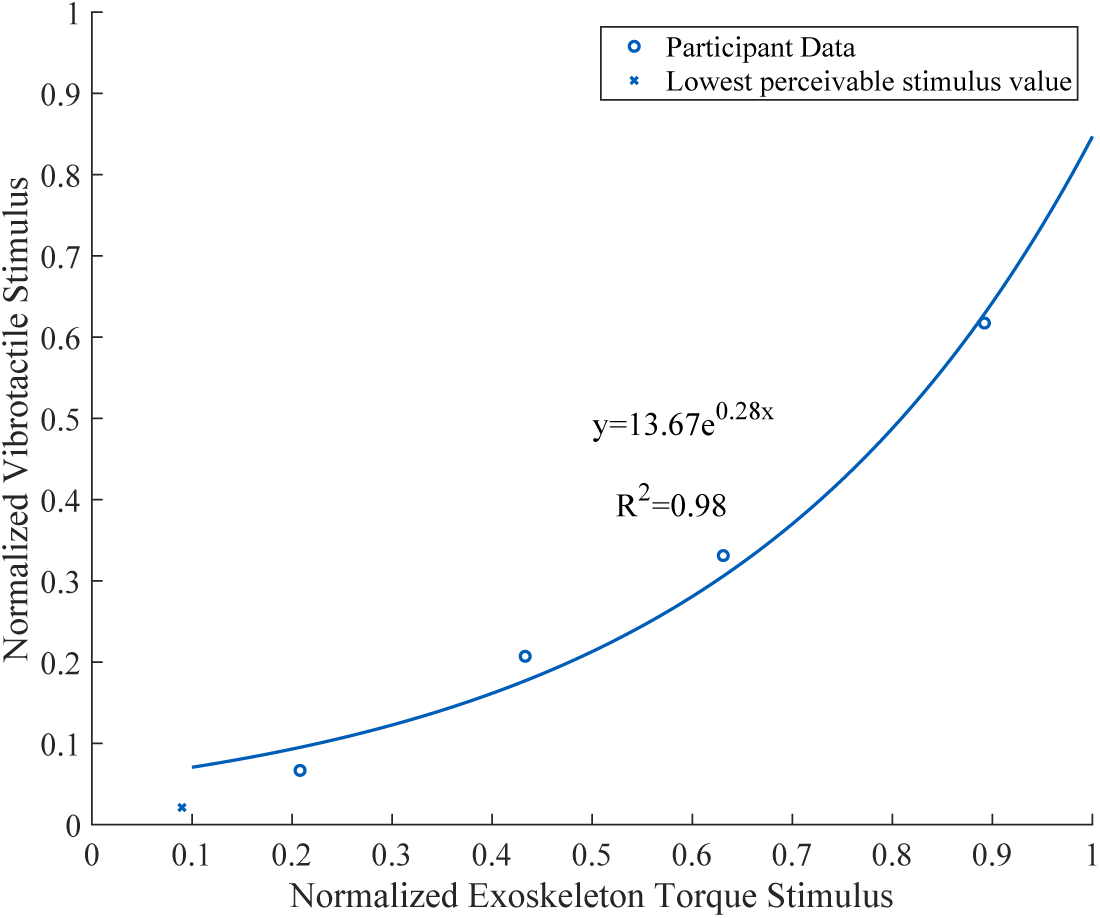
Cross-modal matching curve fit for joint-torque and vibration stimuli for one representative participant. In this example, the normalized vibration stimulus is always matched to a normalized joint-torque stimulus of greater intensity.

### 2.4 Electromyographic Calibration and Processing

While sitting on the stool facing the monitor, one EMG electrode was placed on the participant’s left wrist flexor muscle group. The second EMG electrode was placed on the participant’s left wrist extensor muscle group. The location of these muscle groups was found by asking participants to flex or extend their wrist several times while the experimenter palpated the forearm to find the belly of the appropriate muscle. A ground electrode was placed on the participant’s left elbow. A compression sleeve was placed over the participant’s left arm to secure the electrodes in place. The participant was then instructed to comfortably place their left arm inside the mock prosthesis, which was secured to the participant’s arm using Velcro straps. The base of the prosthesis terminal device was then positioned on top of a support block to achieve the necessary height to perform the experimental task.

The calibration period lasted ten seconds for each muscle group. During the first three seconds, the participant was asked to remain still and maintain a relaxed arm position to gather baseline EMG data. The participant was then asked to perform a series of wrist flexion pulses (seven seconds) followed by a series of wrist extension pulses (seven seconds). Participants were told to keep their contraction effort to normal, everyday levels to mitigate the fatigue associated with the repetitive contractions required to control the prosthesis. The maximum EMG signal for both flexion and extension pulses were averaged and used to normalize the EMG signals for both flexion and extension during the experiment. During prosthesis operation, the raw EMG signal was rectified and smoothed with a 200 ms RMS window. The conditioned signal was then offset adjusted and normalized.

### 2.5 Prosthesis Control

The prosthesis terminal device opening and closing velocity was proportionally controlled by the wrist extension and flexion EMG signals, respectively. Participants used the prosthesis to probe objects, which are described in more detail in Section 2.6.4. During device operation, participants were instructed to close the terminal device to the target aperture range, the lower bound of which is *E*^*t*^. This aperture is measured through the encoder (*E*) on the back of the motor, and was chosen through pilot experiments as the minimum aperture at which a sufficient load cell signal was produced for each of the test objects. *E*^*t*^ corresponds to an aperture of 46.72 mm. When the terminal device is fully open, the aperture is 79.4 mm. During device closing, the prosthesis actuator command was set to zero once the prosthesis aperture reached the closing threshold (*E*^*c*^) which is 103% of *E*^*t*^, and was chosen through pilot experiments to prevent damage to the actuator components and prevent over-gripping the test objects. Even though disabling occurred at 103% of the target aperture, the participant was still able to overshoot past even 110% of *E*^*t*^ due to the momentum of the linear actuator drive. Similarly, during device opening, the prosthesis actuator was disabled once the prosthesis aperture reached the opening threshold (*E*^*o*^) which is 0.7% of *E*^*t*^, and was also chosen through pilot experiments to prevent damage to the actuator components. See Fig. 4a for the signal flow diagram of the prosthesis control.

**Figure 4.**
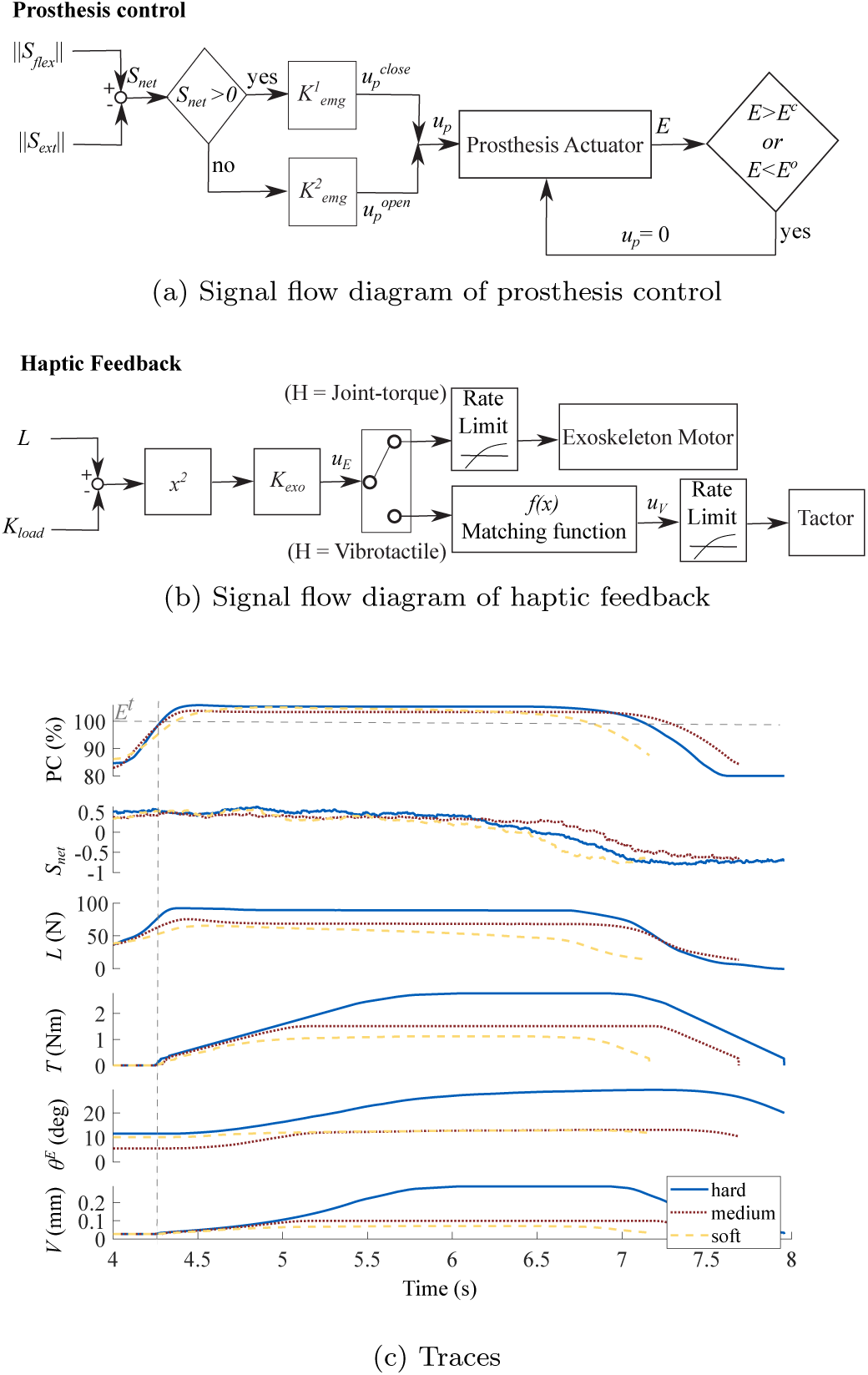
(a) Signal flow diagram of prosthesis control. Terms are described in equations (1) and (2). (b) Signal flow diagram of haptic feedback. Terms are described in equations (3) and (4). The matching function block refers to equation (4). (c) Sample time-series showing the terminal device aperture percent closed (*PC*), the normalized net EMG signal (*S_net_*), the load cell signal (*L*), the exoskeleton output torque (*T*), the exoskeleton elbow angle (*θ*^*E*^), and the vibrotactile skin displacement (*V*) for the hard (solid-blue), medium (red-dot), and soft (yellow-dash) blocks. Traces are truncated to a window after the initial start of object grasp to highlight the rate-limited ramp-up period of the feedback. Note that feedback is only turned on when *PC ≥* 100 (*E*^*t*^, denoted by grey horizontal dashed line. Grey vertical dashed line identifies the time-point when *E*^*t*^ is reached).

The control law for controlling the prosthesis terminal device *u*^*p*^ under proportional EMG control was

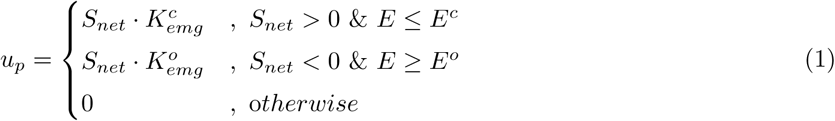

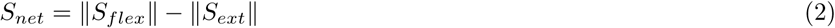

where *S*_*net*_ is the net EMG signal calculated using the normalized EMG wrist flexor signal ‖*S*_*flex*_‖ and the normalized EMG wrist extensor signal ‖*S*_*ext*_‖ as indicated in (2). *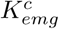* is the proportional gain applied to *S*_*net*_ for device closing, *E* is the encoder reading on the back of the linear actuator motor, and *E*_*c*_ is the encoder threshold during device closing. If the device has closed beyond the threshold (*E > E*^*c*^), or if the participant is producing a larger signal with their extensor muscle (*S*_*net*_ *≤* 0), the motor command for device closing is set to zero. Similarly, 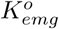 is the gain applied to the difference between EMG flexion and extension signal for opening the terminal device, and *E*_*o*_ is the encoder threshold during device opening. If the terminal device is fully opened (*E < E*^*o*^), or if the participant is producing larger signal with their flexor muscle (*S*_*net*_ *≥* 0), the motor command for device opening is set to zero. In this way, device closing and opening are controlled proportionally with respect to the net EMG signal. Different *K* values for closing and opening the prosthesis were used to account for internal spring in the prosthetic prehensor, which provided a resistive force during closing and an assistive force during opening.

### 2.6 Haptic Feedback Operation

When participants flex their wrist, the EMG flexor signal increases, which in turn drives the prosthesis actuator to pull on the Bowden cable and close the terminal device. A constant offset is subtracted from the load cell signal to account for the reaction force of the terminal device’s internal spring and friction in the cable. Grip force feedback, however, is not provided until the encoder on the back of the motor is greater than or equal to the lower bound of the target range (*E*^*t*^). This encoder (and therefore aperture) threshold provides a measurable and distinct force signal for each of the test objects and was determined through pilot experiments. Once the desired threshold is reached, the load cell signal is held constant to avoid grip force artifacts caused by cable friction, the terminal device spring, and momentum in the cable-drive actuator. Finally, the load cell signal is squared before being sent to the haptic device to provide better separation in the force signals for the test objects. Each haptic device was placed on the contralateral (right arm) of the participant. See Fig. 4b for the signal flow diagram of haptic feedback output.

#### 2.6.1 Joint-Torque Feedback

A gain is applied to the squared net load cell signal and this amplified signal drives the exoskeleton motor. The gain was chosen such that saturation did not occur even when the stiffest block was presented. The output exoskeleton motor command is rate-limited with a rising slew rate of 0.58 Nm/s and a falling slew rate of −1.01 Nm/s to prevent the exoskeleton motor from jerking due to the abrupt signal change that occurs when haptic feedback is turned on, which would be an unintended haptic cue. The exoskeleton was programmed to generate a flexion movement about the user’s elbow. The control law for joint-torque feedback was:

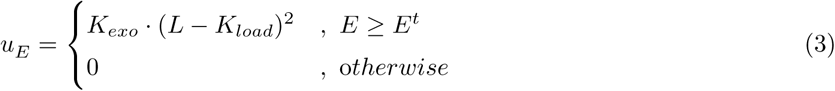

where *K*_*exo*_ is the gain, *L* is the load cell signal, *K*_*load*_ is the constant offset subtracted from the load cell signal, *E* is the encoder reading on the back of the linear actuator motor, and *E*^*t*^ is the target threshold that the terminal device must reach as measured by the encoder. Feedback is initiated once this target aperture is reached. In this way, joint-torque feedback is proportional to the squared load cell signal.

#### 2.6.2 Vibrotactile Feedback

A gain is applied to the squared net load cell signal. The output exoskeleton motor command is mapped to a vibration command using the curve fit parameters identified during cross-modal matching. The output is also rate-limited with a rising slew rate of 1 mm/s and a falling slew rate of 1.2 mm/s, although perception of the cue was discrete. The control law for vibrotactile feedback was:

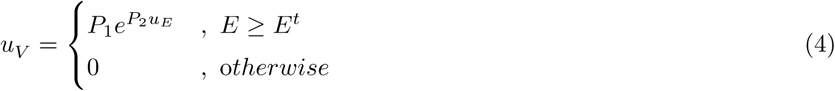

where *P*_1_ is the first coefficient of the two parameter exponential function found during the cross-modal matching, *P*_2_ is the second coefficient of this function, and *u*^*E*^, *E*, and *E*^*t*^ are as described in (3). In this way, vibrotactile feedback is presented exponentially with respect to joint-torque feedback according to the cross-modal matching model fit.

#### 2.6.3 Sample Time-Series

The system operation can be visualized in Fig. 4c for the soft, medium, and hard blocks. A positive EMG signal drives the motor to close the prosthesis terminal device around the test block. After reaching the threshold aperture (*E*^*t*^), feedback is initiated. A negative EMG signal drives the motor in the opposite direction to open the prosthesis terminal device, causing a decrease in the haptic stimulus intensity.

#### 2.6.4 Stimuli

Participants were asked to discriminate pairs of blocks with different stiffnesses in three different conditions: no haptic feedback, vibrotactile feedback, and joint-torque feedback. There were four Smooth-On silicon blocks: soft (Ecoflex 20; shore hardness 00-20), medium (Ecoflex 30; shore hardness 00-30), hard (Dragon Skin 10; shore hardness 10A), and extra-hard (Dragon Skin 20; shore hardness 20A). The extra-hard block was used only for catch trials. Participants also interacted with two sample blocks during training, Ecoflex 50 (shore hardness 00-50) and SORTA-Clear 40 (shore hardness 40A), which were not used during the actual experiment. Each block measures 67 mm by 30 mm by 142 mm. Blocks were fit into custom 3D-printed holders, as shown in Fig. 5.

**Figure 5.**
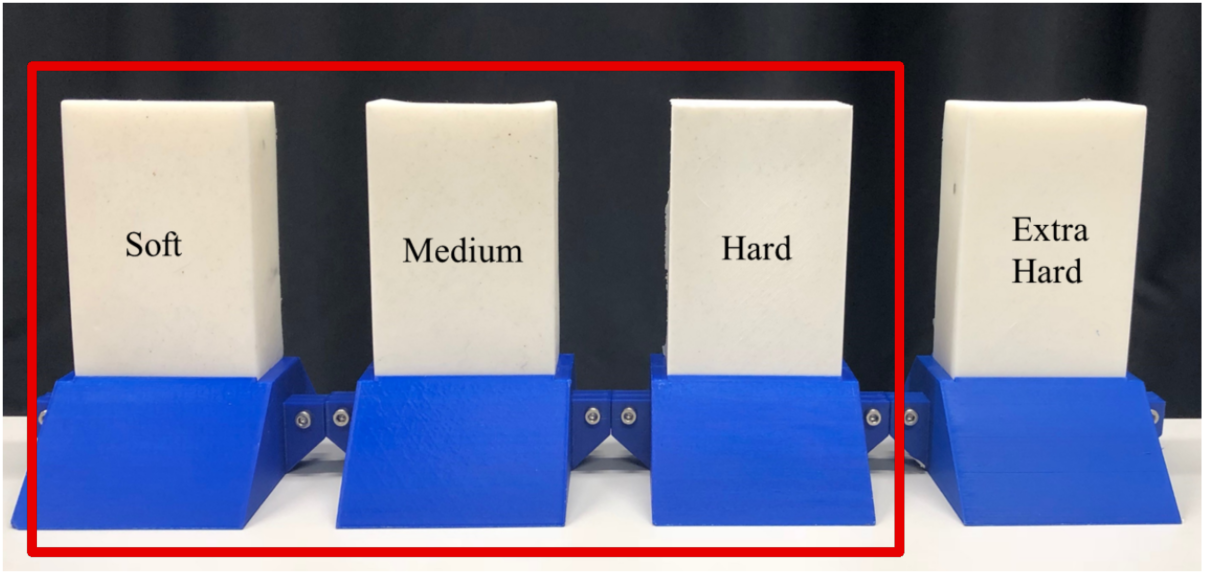
Test blocks used for the object discrimination task, arranged in their 3D-printed holders from least to most stiff (left to right). Only data from the outlined soft, medium, and hard blocks were processed, while the extra hard block was used only for catch trials.

### 2.7 Protocol

Participants were split into two groups (A and B) in an alternating fashion. Group A performed the task in the following condition order: (1) no feedback, (2) joint-torque feedback, and (3) vibrotactile feedback. Group B performed the task in the following condition order: (1) no feedback, (2) vibrotactile feedback, and (3) joint-torque feedback. After completing the cross-modal matching, participants completed a brief survey regarding their demographics, handedness, and their perceived ability to successfully complete the cross-modal matching experiment. After completing each condition, subjects completed a post-condition survey. The survey was a mix of short-answer and slider-scale qualitative questions about participants’ perceived performance and evaluation of the task.

Prior to starting the experiment, participants were allowed to probe the sample blocks with their free hand to understand how the reaction force of the object is related to its stiffness. These sample blocks were then placed between the jaws of the terminal device on the prosthesis. Vision of the blocks was occluded by the computer monitor and two pieces of polyester fabric; participants were only able to observe the monitor in front of them which displayed a graph indicating the percentage closed of the terminal device. Participants were asked to flex their wrist to close the terminal device and aim for between 100% and 110% of the target threshold, *E*^*t*^, which corresponds to a range of 46.72 to 38.11 mm terminal device aperture. The upper bound was chosen through pilot experiments as the minimum amount of overshoot over *E*^*t*^ that our novice participants could consistently achieve with myoelectric control, while also providing enough of a range for noticeable visual differences between blocks in a pair. This upper bound also encourages the participants to maintain consistency across all trials. Participants were allowed to practice on both of the sample blocks while the experimenter explained which block was stiffer. Sample blocks were not used in the actual test. See Fig. 6 for experiment setup.

**Figure 6.**
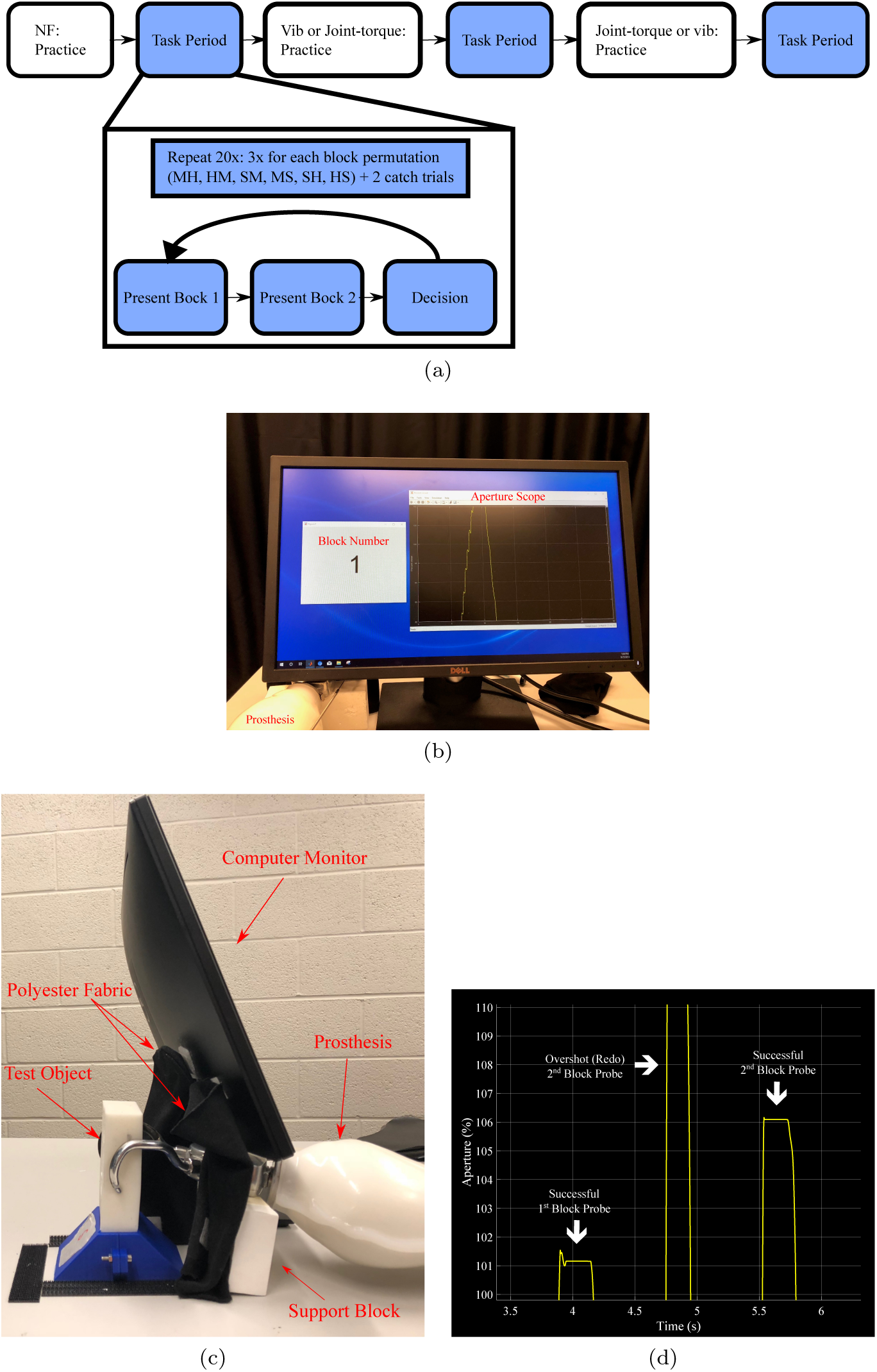
(a) Experiment procedure flow diagram, where NF indicates no feedback; H, M, and S refer to hard, medium, and soft blocks (b) View of experimental setup from the participant’s point of view, (c) Sideview of the experimental setup. Fabric and computer monitor occlude the participant’s view of the block. (d) Example probes – participant needs to reach between 100% and 110% of the target aperture *E*^*t*^. If they undershoot or overshoot this target, they are required to try again.

After the participant was comfortable with the prosthesis control, the object discrimination experiment began. The three test blocks (soft, medium, hard) were presented in six pairwise permutations whose order was randomized. Each pairwise permutation was repeated three times for a total of 18 presentations. Two catch trials were randomly placed throughout the 18 trials, making 20 trials total. The catch trial always consisted of a soft and extra-hard block comparison. If the participant failed to accurately identify the extra-hard block as being the stiffer block in both catch trials, for either haptic feedback condition, the study was terminated. Catch trials, regardless of whether they were successful or not, were not included in the data analysis, but were only used to indicate if participants were paying adequate attention to the feedback, or to identify potential issues with the experimental device operation. During each trial, the participant was allowed a single probe of each block. If the participant’s *S*_*net*_ command caused the terminal device aperture to overshoot or undershoot the target aperture range, participants were instructed to repeat the probe. In addition to the aperture display, the monitor also displayed prompts for block 1, block 2, or probe redo (See Fig. 6). Participants were able to repeat a single probe of each block within a pair as many times as they desired. After probing both blocks to satisfaction, participants verbally indicated which block they perceived to be stiffer, and the experimenter recorded this answer on the computer.

### 2.8 Metrics and Statistical Analysis

The primary performance metric was the percent accuracy for object pair discrimination. Logistic mixed-effects models were implemented in R version 3.4.3 to compare accuracy between block combinations and feedback conditions. Within the model, the interaction of block combination and condition was a fixed effect.

As a secondary performance assessment, we evaluated how participants’ choices in the object discrimination task related to various incidental cues associated with task execution. These cues were quantified by a set of features that were calculated for every block presented to the participant and are described below. Features were chosen such that the larger the value, the higher the presumed stiffness of the block. These features include grip aperture, probe difficulty, amount of overshoot and undershoot of the target range, amount of EMG activity, and stimulus intensity for each haptic condition. In the second model, the interaction of aperture and condition, interaction of probe difficulty and condition, three-way interaction of probe difficulty, aperture, and condition, and interaction of stimulus intensity and condition were fixed effects.

Lastly, we used a logistic mixed model to determine if the strategy (fixed effect) used in the joint-torque condition affected the accuracy outcome.

All fixed-effects for each of the models were chosen based on the lowest Bayesian Information Criterion. Participants were included as random effects in all models. Multiple comparisons were accounted for using Bonferroni adjusted p-values.

#### 2.8.1 Probing Difficulty

The probing difficulty for each block, *PD*, quantifies how often participants either undershot or overshot the target aperture range (100%-110% of *E*^*t*^) when probing the block. Participants were allowed multiple probes of a block if and only if they overshot or undershot the target range, or if they asked to repeat the block pair. The probing difficulty is defined as:

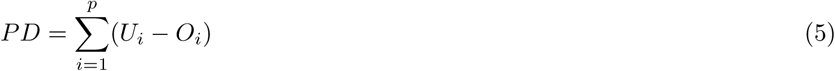

where *p* is the number of probes for a given block, *U*^*i*^ is a binary variable indicating whether the target range was undershot for the *i*th probe, and *O*^*i*^ is a binary variable indicating whether the target range was overshot for the *i*th probe. With the prosthesis, participants were more likely to undershoot the stiffer blocks and overshoot the softer blocks. Essentially, *PD* compares the number of times the block is undershot to how many times it is overshot. Therefore, as *PD* increases, so too does the presumed likelihood of the block being stiff.

#### 2.8.2 Aperture

The aperture feature for a given block, *A*, quantifies the average normalized value of the prosthesis aperture inside the target range, and is defined as:

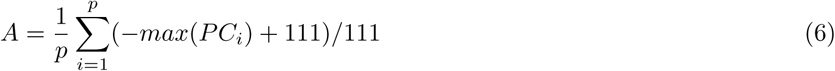

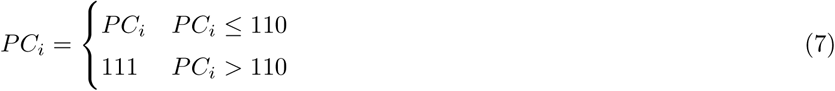

where *p* is the total number of probes for a given block, and *PC*_*i*_ is the maximum percent closed of the terminal device displayed in the scope for the *i*th probe. Since the maximum value on the scope that participants are able to view is 110%, any aperture above this value appears visually the same to the participant. Therefore, *PC*_*i*_ is adjusted according to (7) such that any aperture above 110% is represented the same. Essentially, (6) converts the terminal device’s final aperture for each block into an associated stiffness value. The participant is likely to reach a higher aperture for softer blocks, and lower apertures for stiffer blocks. Using (6), we invert the relationship between stiffness and block deformation, so that as *A* increases, so too does the presumed likelihood of the block being stiff.

#### 2.8.3 Net EMG

The net EMG feature for each block, *S* quantifies the average muscle activity for a given block and is defined as:

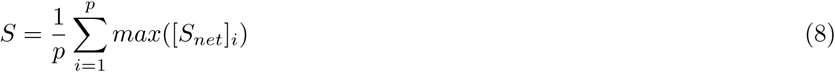

where *p* is the total number of probes and [*S*_*net*_]_*i*_ is the net EMG signal as described in (2) for the *i*th probe. The EMG value is related to the amount of muscle activity; participants are more likely to generate more wrist flexor muscle activity to reach the target range for stiffer blocks, and less so for softer blocks. Therefore, as *S* increases, so too does the presumed likelihood of the block being stiff.

#### 2.8.4 Joint-Torque Stimulus Intensity

The joint-torque stimulus feature for each block, 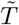, quantifies the average normalized stimulus intensity of the joint-torque feedback for a given block and is defined as:

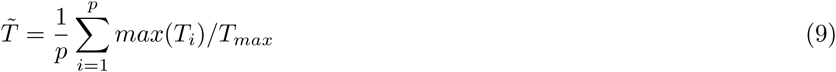

where *p* is the number of probes for the block and *T*_*i*_ is the joint-torque signal for the *i*th probe. *T*_*i*_ was normalized by the maximum joint-torque stimulus, *T*_*max*_, or in this case, 3.23 Nm. Participants are likely to receive a larger joint-torque stimulus for a stiffer block than for a softer block. Therefore, as *T* increases, so too does the presumed likelihood of the block being stiff.

#### 2.8.5 Vibrotactile Stimulus Intensity

The vibrotactile feedback feature, 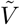, quantifies the average normalized vibrotactile stimulus intensity for a given block and is defined as:

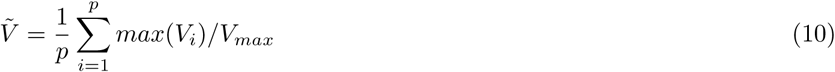

where *p* is the total number of probes for that block, *V*_*i*_ is the vibration amplitude for the *i*th probe. *V*_*max*_ is the maximum possible amplitude for a particular participant according to their cross-modal matching curve, up to the C-2 Tactor hardware limits (0.499 mm displacement). Participants are likely to receive a larger vibrotactile stimulus for a stiffer block than for a softer block. Therefore, as 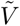 increases, so too does the presumed likelihood of the block being stiff.

### 2.9 Block Pair Feature Comparison

Once all the above features have been calculated for both blocks in a pair, the first block’s set of features is subtracted from the second block’s set of features. The resulting difference indicates the presumed difference in stiffness between the two blocks; if the value is positive, then the presumed stiffness of the second block is larger than that of the first block, however, if the value is negative, the presumed stiffness of the first block is larger than that of the second block.

### 2.10 Survey

After each condition, participants were asked to complete a survey consisting of questions related to their perceived performance, and subjective assessments of the tasks and conditions. These questions are based on the NASA TLX questionnaire. The questions were a mix of short answer and sliding-scale questions on a rating scale of 0-100. Only the rating questions will be discussed further. The first question asked participants to rank how physically demanding the condition was and the second question asked participants to rank how mentally demanding the condition was. The third question asked how hurried or rushed the task was, while the fourth question asked participants to rate their perceived accuracy in each condition. Questions six and seven asked participants to rate how hard they had to work to achieve their level of performance and how discouraged or frustrated they were during the task. The eighth question asked how useful the participants thought each of the feedback types were. The final question recorded what method they used to discriminate torque.

## 3. Results

In analyzing the data, we excluded the data from two participants. The first participant made multiple probes of each block even after reaching the target range, despite repeated instruction to not do so. Their results are inconsistent with the other participants given that this participant received more information per trial from which to discriminate the objects. This was inconsistent with the experimenter’s instructions to probe each block a single time, unless they undershot or overshot the target range. The second participant failed each of the catch trials during the joint-torque condition, likely due to a hardware issue that caused the load cell signal to appear the same for many block pairs. This issue was fixed after this participant’s session. Our analysis will therefore focus on the ten remaining participants, of which seven were male, three were female, and the average age was 24 years. Five participants were in group A, and five were in group B.

### 3.1 Accuracy

In determining the best model, it was found that both trial and participant group were not significant fixed effects, and additionally increased the Bayesian information criterion. Therefore, these effects were not included in the final chosen model (see Section 2.8). Accuracy was better in the vibrotactile feedback than the no feedback condition for the medium-hard block combination. Accuracy was also significantly better in both the vibrotactile and joint-torque feedback conditions than the no feedback condition for the soft-hard block combination. There was no difference between any conditions in the soft-medium block combination, and no differences between the vibrotactile and joint-torque conditions in any block combination. Other comparisons were not analyzed. As the interactions were significant, main effects were not analyzed, as recommended in [25]. The results are summarized in Table 1 and displayed in Fig. 7.

**Table 1.**
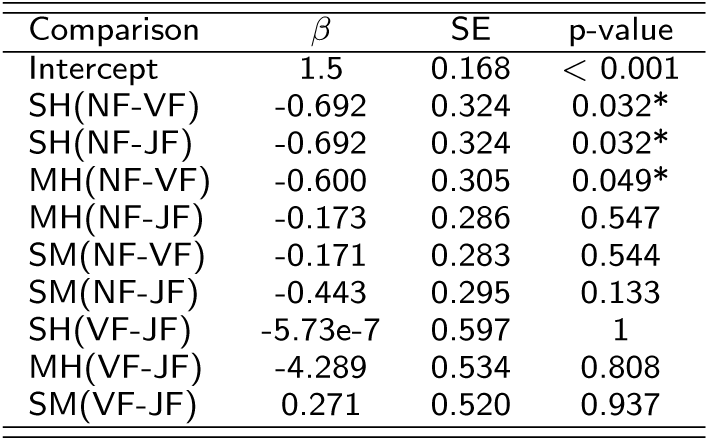
Main model fixed effect results. NF is no haptic feedback, VF is vibrotactile feedback, and JF is joint-torque feedback. MH, SM, and SH indicate medium-hard, soft-medium, and soft-hard block combinations, respectively. * indicates p *<* 0.05.

| Comparison | $\beta$ | SE | p-value |
| --- | --- | --- | --- |
| Intercept | 1.5 | 0.168 | < 0.001 |
| SH(NF-VF) | -0.692 | 0.324 | 0.032* |
| SH(NF-JF) | -0.692 | 0.324 | 0.032* |
| MH(NF-VF) | -0.600 | 0.305 | 0.049* |
| MH(NF-JF) | -0.173 | 0.286 | 0.547 |
| SM(NF-VF) | -0.171 | 0.283 | 0.544 |
| SM(NF-JF) | -0.443 | 0.295 | 0.133 |
| SH(VF-JF) | -5.73e-7 | 0.597 | 1 |
| MH(VF-JF) | -4.289 | 0.534 | 0.808 |
| SM(VF-JF) | 0.271 | 0.520 | 0.937 |

**Figure 7.**
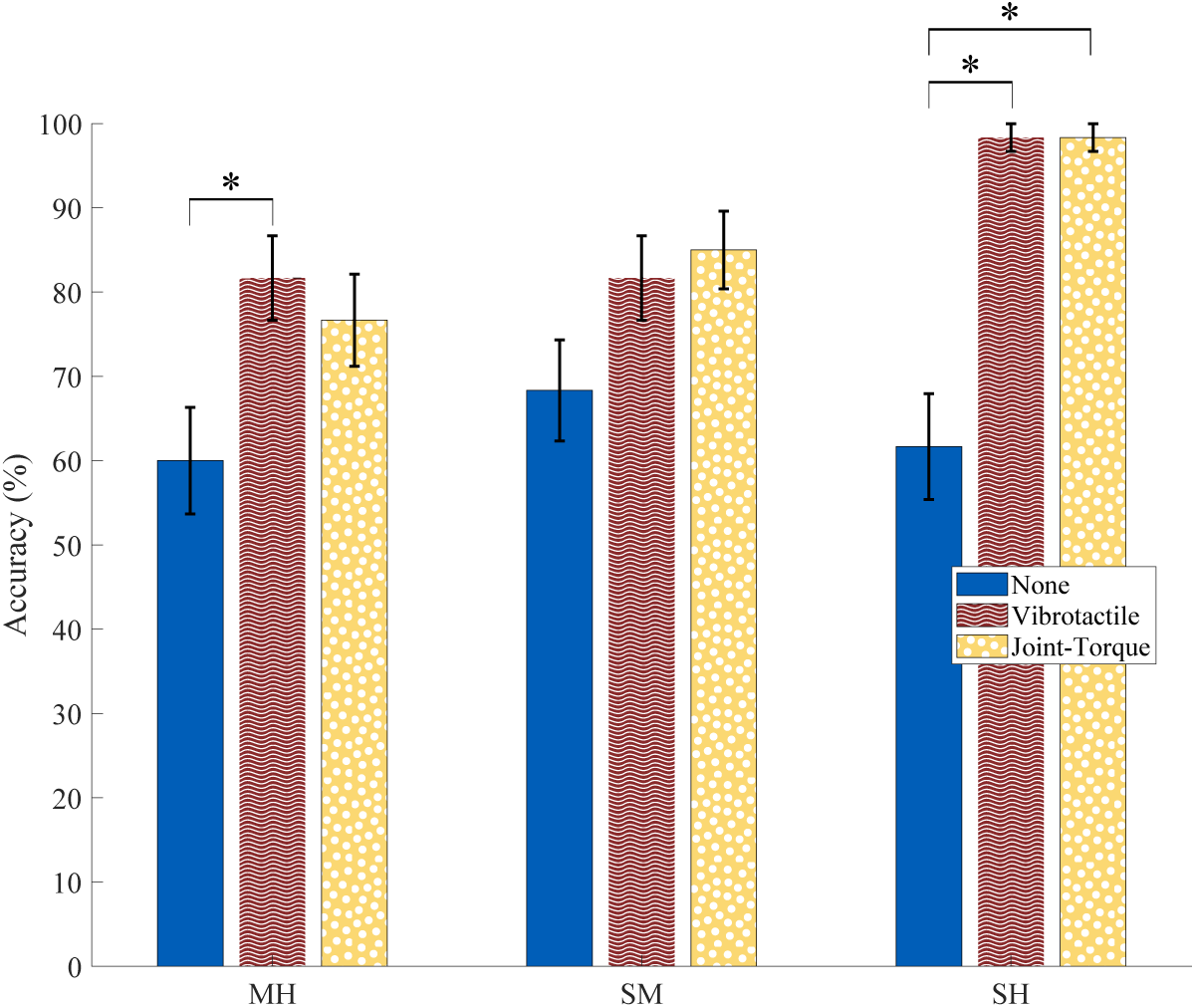
Average accuracy for block combination (MH: medium-hard, SM: soft-medium, SH: soft-hard) and task condition. Error bars represent 1 standard deviation. * indicates p *<* 0.05 and ** indicates p *<* 0.01

### 3.2 Feature Analysis

In this section, we present an analysis of what features were likely important when participants were deciding which block was stiffer. Features like probing difficulty, aperture, and EMG depended mostly on the participant’s ability to control the prosthesis. Haptic feedback features are less related to the participant’s ability to control the terminal device and more related to the physical properties of the block. In determining the optimal fixed effects to use, it was found that EMG was not significant, and therefore are not included in the final chosen model (see Section 2.8).

Results are summarized in Table 2. Participants in the no feedback condition were more likely to choose the second block in the pair as being more stiff when the difference in aperture between the second and first block was more positive. Likewise, participants in the joint-torque condition were more likely to choose the second block in the pair as being more stiff when the difference in aperture between the second and first block was more positive. Participants in all three conditions were more likely to choose the second block as stiffer when the difference in probe difficulty between the second and first block was more positive. Participants in the two haptic feedback conditions were more likely to choose the second block as stiffer when the difference in the respective haptic stimulus intensities between the second and first block was more positive. Only in the no feedback condition were participants more likely to choose the second block as stiffer when both the difference in aperture and probe difficulty between the second and first block were more positive. The intercept was also significant (*β* = 0.33, *SE* = 0.12, p = 0.006). The actual response rate for choosing the second block is shown in Fig. 8, along with the above model fit to the actual data to depict the probability of choosing the second block given each feature.

**Table 2.** Feature fixed effect results. NF is no haptic feedback, VF signifies vibrotactile feedback, and JF signifies joint-torque feedback. A, PD, and SI refer to the aperture, probe difficulty, and stimulus intensity features, respectively. * indicates p *<* 0.05, ** indicates p *<* 0.01, and *** indicates p*<* 0.001

| Comparison | $\beta$ | SE | p-value |
| --- | --- | --- | --- |
| Intercept | 0.329 | 0.120 | 0.006** |
| A:NF | 1.52 | 0.493 | 0.002** |
| A:VF | 0.190 | 0.590 | 0.747 |
| A:JF | 1.08 | 0.537 | 0.043* |
| PD:NF | 0.421 | 0.104 | <0.001*** |
| PD:VF | 0.327 | 0.153 | 0.033* |
| PD:JF | 0.326 | 0.140 | 0.019* |
| A:PD:NF | 0.493 | 0.237 | 0.038* |
| A:PD:VF | 0.064 | 0.361 | 0.860 |
| A:PD:JF | -0.438 | 0.304 | 0.149 |
| SI:VF | 5.04 | 0.794 | <0.001*** |
| SI:JF | 6.58 | 0.995 | <0.001*** |

**Figure 8.**
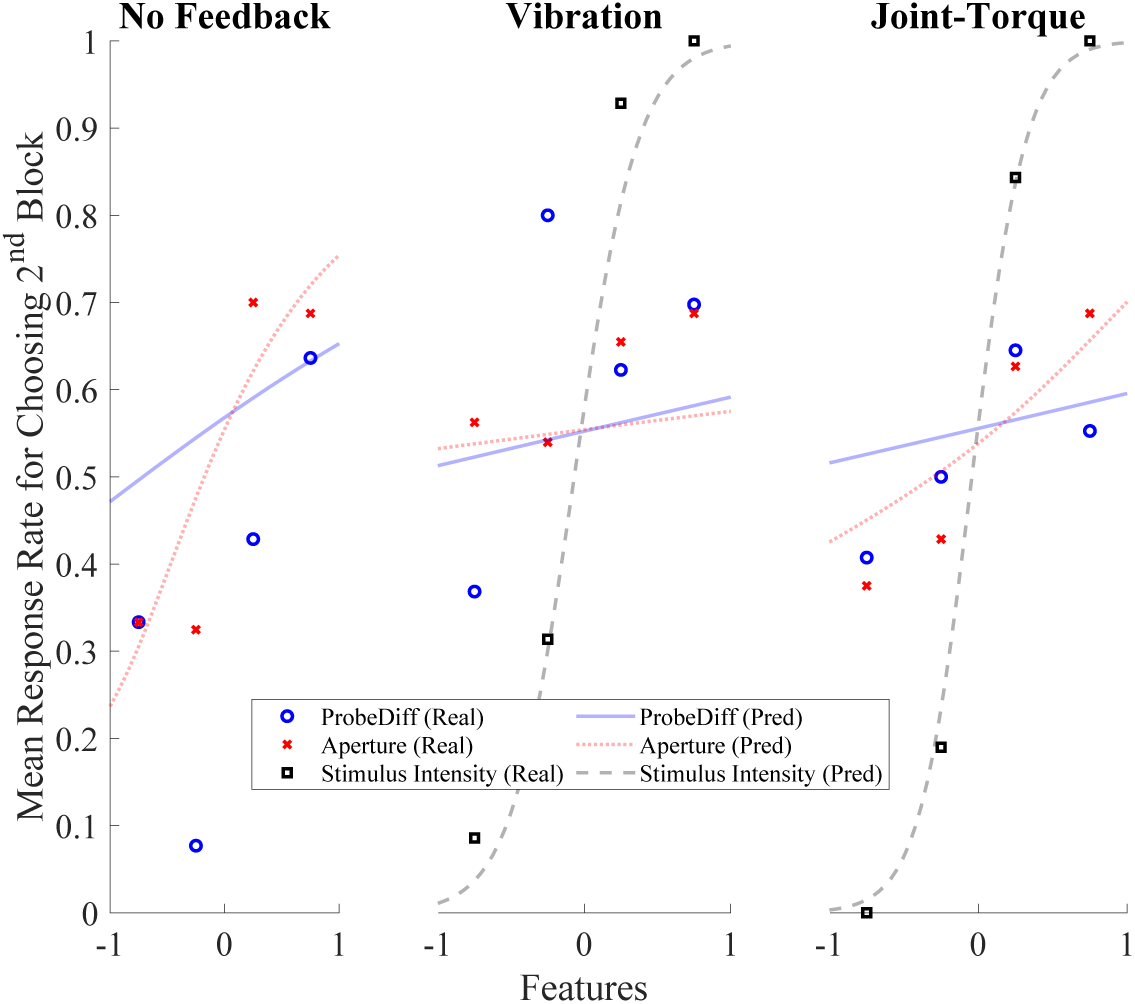
The actual response rate of choosing the second block given the Aperture, Probe Difficulty and Stimulus Intensity features. Plotted alongside the real data is the model’s prediction of response rate for these features. Features are normalized between −1 and 1. The higher the feature value, the bigger the difference was for that feature between the first block and the second block. If the feature was negative, the first block feature was larger than the second block.

### 3.3 Surveys

Surveys indicated significant differences between each haptic condition and the no feedback condition in all rating-style questions except for physical effort. However, there were no significant differences between joint-torque and vibrotactile feedback in any of the questions.

Participants perceived themselves to be significantly more accurate with vibrotactile (p *<* 0.001) and joint-torque feedback (p *<* 0.001) feedback than no feedback. Participants felt that the vibrotactile (p = 0.006) and joint-torque (p = 0.018) feedback were significantly more useful in the task than no feedback. Participants felt that their performance in the task required significantly less effort with vibrotactile (p *<* 0.001) and joint-torque (p = 0.002) feedback than with no feedback. Participants felt less of a mental demand with the vibrotactile (p *<* 0.001) and joint-torque (p *<* 0.001) feedback than with no feedback. Finally, participants felt the vibrotactile (p *<* 0.001) and joint-torque (p *<* 0.001) conditions were significantly less frustrating than the no feedback.

Six participants said they allowed the exoskeleton to move their arm, while three participants actively moved their arm to feel the torque. One participant noted that they passively allowed elbow flexion for large torques, but actively moved their arm back and forth for small torques. There was no effect of strategy in the joint-torque condition on the accuracy outcome, even when removing the participant who used a combination of strategies. There was a significant effect of intercept (*β* = 1.87, SE = 0.269, p *<* 0.001).

## 4. Discussion

In this study, we investigated the utility of cutaneous and kinesthetic haptic feedback in a myoelectric prosthesis and found that both feedback modalities resulted in performance that was significantly better than the clinical standard, no haptic feedback. We arrived at this conclusion through an experimental investigation involving a custom myoelectric prosthesis that can be worn by non-amputee participants and features joint-torque (kinesthetic) and vibrotactile (cutaneous) feedback that can be conditionally removed. In addition, we employed a cross-modality matching technique that generated user-defined equivalence mappings between joint-torque and vibrotactile stimulus intensities. In this way, we are able to compare each feedback modality to the clinical standard, as well as directly compare the two different modalities against each other without the confound caused by perceptual differences due to stimulus intensity.

### 4.1 Task Performance

The current research on haptic feedback in upper extremity prostheses is mixed with regard to the feedback’s potential utility. Not all investigations of haptic feedback in prostheses have resulted in improved functionality. In previous work done by members of our research group, it was found that neither vibrotactile nor joint-torque feedback improved grasp and lift performance over vision with a myoelectric prosthesis [16]. Likewise, Saunders *et al* found that vibrotactile feedback was only useful in a grasp and lift task with a myoelectric prosthesis when feedforward uncertainty was present in the control loop [26]. On the contrary, it has previously been shown that vibrotactile feedback prevents amputees from breaking objects when wearing a myoelectric prosthesis [12]. Likewise, a previous study done by members of our lab showed that joint-torque feedback provided added utility over vision in a stiffness discrimination task with a body-powered prosthesis [7]. Other studies have also shown that stiffness discrimination is improved with the addition of haptic feedback [27], [28]. Therefore, our finding, that the addition of vibrotactile and joint-torque feedback to myoelectric prostheses allows for improved object discrimination over no feedback for the soft-hard block combination, supports the argument that both cutaneous and kinesthetic forms of feedback can provide added utility and function to a myoelectric prosthesis. We believe the decreased performance in the no haptic condition was likely do to the absence of grip force information and not due to learning effects as we found no significant effect of trial order in our original model selection. Although the efficacy of haptic feedback appears to be related to how close the blocks are in stiffness, incorporating haptic feedback into a prosthesis would give users a better chance at object discrimination, which is critical for many activities of daily living.

Our secondary result indicated that there was no difference in object discrimination between joint-torque feedback and vibrotactile feedback in any of the block combinations. The literature comparing joint-torque feedback and vibrotactile feedback is limited to research done previously by members of this research group. Brown *et al* found that there was no difference between joint-torque and vibrotactile feedback in a grasp and lift task using a myoelectric prosthesis [16]. Similarly, Brown *et al* found that there was no difference in kinesthetic feedback delivered to a different part of the body than the part used for exploration (non-colocated) and vibrotactile feedback in a single-DoF spring stiffness discrimination task [29].

More broadly, studies have explored differences between kinesthetic and cutaneous feedback. In an angle discrimination task, Frisoli *et al* found no difference between kinesthetic and cutaneous feedback [30]. Recently, Kamikawa and Okamura found that hand-grounded kinesthetic feedback performed better than skin deformation (cutaneous) feedback in a force-matching task relative to a world-grounded kinesthetic device [31]. This result, while contradictory to our own, might well be explained by the experimental task used, the kinesthetic control condition employed, or the colocated feedback (action originates from the same body part as where the haptic feedback is felt), which was shown previously by members of our research group to be superior to cutaneous feedback [29].

Any stimulus can be reduced to its intensity and somatosensory encoding. In doing cross-modal matching, we attempted to equalize the intensities for each modality to compare differences in somatosensory encoding only. The lack of significant differences between the joint-torque and vibrotactile conditions indicate that participants’ ability to encode force as vibration or torque was not different, as both modalities were sensory substitutions of force. If one of the modalities was not a sensory substitution, it is possible that differences between modalities would emerge, as in [31].

It is also possible that friction in the exoskeleton masked the object’s true force, thereby making it difficult to discriminate objects with relatively close stiffness values. For both vibrotactile and joint-torque conditions, we calculated the difference in haptic feedback stimulus intensity for each block pair and determined how many were below the just noticeable difference (JND). We found that 30%, 3.3%, and 0% of the stimuli differences were below an elbow joint-torque JND of 0.13 as reported in [23] for the medium-hard, soft-medium, and soft-hard block pairs, respectively. For vibrotacile difference thresholds, Rothenberg *et al* determined that the JND for forearm stimulation at 250 Hz was 0.2, while Mahns *et al* found that it was 0.3 [24]. Therefore, we chose a difference threshold of 0.25 which is in between these reported values. The proportion of stimulus intensity differences within a block pair less than or equal to 0.25 was 5%, 8.3%, and 0% for the medium-hard, soft-medium, and soft-hard block pairs, respectively. Compared to the joint-torque condition, the vibrotactile condition had much fewer pair differences below the JND in the medium-hard block pair combination, which may have contributed to it being significantly better than no haptic feedback, whereas joint-torque was not. In addition to improving the proportion of differences above the JND, an increased number of participants may also have helped to strengthen the significance of the results in the other combinations and conditions.

### 4.2 Feature Analysis

In addition to analyzing performance differences between the task conditions, we also examined how participants chose which block was stiffer. In all conditions, participants were privy to a number of incidental cues that could be used to make an educated guess as to which block was stiffer. Those cues included visual feedback of the terminal device aperture, visual indications of the overshoot and undershoot of the target aperture, and for the haptic feedback conditions, the perceived haptic stimulus intensity. Our analysis found that, in the absence of haptic feedback, participants’ decisions as to which block was stiffer aligned well with the visual based indicators, even though they did not always correlate with the true object stiffness. When feedback, kinesthetic or cutaneous, was available, reliance on these incidental cues diminished. This reliance on incidental cues is similar to the behavior anecdotally observed in amputees who wear myoelectric prostheses and must rely on visual feedback to properly control and understand interactions between the prosthesis and the environment. In a study done by Subah *et al*, it was found that in a reach-and-grasp task with a prosthesis simulator, both able-bodied users and amputees extensively monitored their virtual hand during grasping [32]. This is in direct contrast with able-bodied individuals using their intact hand to manipulate objects, as their gaze is rarely fixated on the hand, but rather on the object [33]. By adding haptic feedback, it is possible to reduce dependence on visual cues, thereby allowing amputees to focus attention elsewhere. In fact, Raveh *et al* showed that the addition of vibrotactile feedback in a myoelectric prosthesis decreased the time to complete object manipulation tasks alongside a secondary task, indicating that feedback reduced some of the cognitive load associated with the prosthesis task [10].

Interestingly, in the joint-torque feedback condition, in addition to the torque intensity, the aperture cue was also significant for the participants’ choice, while the exoskeleton angle was not. Previously it was noted that 30% of stimuli pairs in the medium-hard block combination in the joint-torque condition were below the JND. This might have led participants to rely more on visual cues like aperture when the feedback was hard to discern.

### 4.3 Surveys

The subjective survey results support the quantitative differences found in task performance between the haptic feedback conditions and the no feedback condition. Participants also rated that both haptic conditions were less mentally taxing and required less effort than the no haptic feedback condition, which supports our feature analysis results. Our statistical analysis indicated that there was no difference in which method participants used to feel the joint-torque; however, it may be worth running a future full experiment on whether passive or active resistance has a significant effect on accuracy.

### 4.4 Limitations

It should be noted, that while our current results have positive implications for prosthesis development, there were several study limitations that should be addressed in future empirical investigations.

First, all haptic feedback was provided to the contralateral arm, thereby making it less natural than providing ipsilateral feedback. Chatterjee *et al* along with investigations from members of our research group (Brown *et al*) showed that haptic feedback on the contralateral limb had reduced performance compared to haptic feedback on the ipsilateral limb [29, 34]. In this present study, we were limited by the exoskeleton, as it was infeasible to put this device on the same arm as the prosthesis. Despite this limitation, we were still able to show significant improvement over no haptic feedback.

Second, the no haptic feedback condition substituted pure visual feedback of the block deformations with a display scope of aperture. Therefore, it was an interpreted method of visual feedback, which may or may not have been more difficult than pure visual feedback. As it was possible to overshoot on any of the blocks, especially for able-bodied individuals who are not experts at EMG control, aperture is not a wholly reliable indicator of stiffness for this particular setup. Therefore, actually seeing the blocks deform may have improved accuracy. On the other hand, participants would not receive quantifiable visual measurements without the scope. We initially decided to use the scope method to ensure participants were consistent in their grip aperture, since closing down more on a soft object could yield a similar grip force to closing down less on a hard object. Additionally, this method allowed us to investigate how much participants used visual or haptic feedback cues in making their decision. A future alternative approach to visual feedback would be to limit the closure of the prosthesis to a set amount and provide feedback on grip force that is discounted by the aperture. Therefore, differences in grip aperture between block pairs would not affect the stimulus feedback.

Third, the load cell was at times sensitive to the relative placement of the prosthesis. The orientation and position of the participant’s arm controlled the slack in the Bowden cable, which in turn controlled the friction in the linear actuator. The orientation could change as the participant made small shifts throughout the duration of the experiment. As the load cell measures the tension in the cable, these values could become inconsistent between two test blocks. We attempted to remedy this issue by marking the places on the table where the prosthesis should rest, and on each of the blocks where the terminal device should make contact. In the future, the interior of the Bowden cable could be lined with Teflon to reduce friction. Although it was possible to simplify the setup by detaching the prosthesis from the subject completely, we wanted to encourage embodiment, and therefore attempted to mimic the amputee’s physical experience as closely as possible.

Fourth, the haptic feedback was held constant after reaching the target aperture threshold (*E*^*t*^), which is not representative of true force perception when squeezing objects. We avoided the natural increase in force that occurs when squeezing an object because the load cell, in addition to measuring the object force, also measured the spring force of the terminal device and the friction in the Bowden cable, which could mask the force of the block and cause difficulty in perceiving the difference between two blocks. By holding the feedback constant after passing *E*^*t*^, we intended to ensure distinct feedback for each test object despite the hardware limitations. This limitation, in addition to the ones mentioned above, would not come into play in a traditional myoelectric prosthesis or a body-powered device as there would be no linear actuator or load cell. Here, we attempted to control for these factors. While cable friction is present in standard body-powered devices, amputees learn to account for this friction with long-term repeated use. We were unable to provide the amount of training for our able-bodied participants to reach this level of proficiency. Additionally, even though the zero-order hold on feedback reduces to force discrimination, the inclusion of visual information encourages the integration of both force and aperture. Consequently, it is feasible for participants to discriminate the various objects based on more than just the haptic feedback of force. Finally, it is worth mentioning that a commercial myoelectric prosthesis outfitted with force sensors on the end effector could be used in future experiments. As a final limitation, we tested only a few able-bodied individuals. In future studies, amputees will be evaluated as their performance may differ in the no haptic feedback condition, given their prior experience with myoelectric prostheses. In addition, we will also use maximum voluntary contraction (MVC) to calibrate EMG control. As our feature analysis is a model, we cannot draw definitive conclusions on how participants actually used the cues, and whether the addition of haptic feedback did indeed reduce cognitive loading. In future experiments, direct measures of cognitive load should be included.

Previous literature has shown that myoelectric prostheses may benefit from the addition of haptic feedback [7,18,21]. In this study, we show that task performance is equally improved over no haptic feedback, regardless of the specific haptic feedback modality used. Contrary to our hypothesis, kinesthetic feedback was not superior to cutaneous feedback; perhaps one reason is that the dynamics of EMG are not exactly coupled to the myoelectric terminal device like the dynamics of the body are coupled to body-powered terminal device. In some respects, the mechanical operation of the body-powered prosthesis provides a colocated form of kinesthetic feedback, which as shown in our previous research, is superior to non-colocated cutaneous and kinesthetic feedback [29]. It may be worth investigating whether there are performance differences between colocated (close to the EMG sensors) and non-colocated (far from the EMG sensors) feedback in a myoelectric prosthesis, as well as the effect of long-term training on the efficacy of haptic feedback. As it stands currently, cutaneous sensory substitution, which is cheaper and simpler to implement, will likely improve object discrimination performance over no haptic feedback.

## 5 Conclusion

In this study, we compared the effects of joint-torque and vibrotactile feedback in an object discrimination task using a myoelectric prosthesis. Prior to experimentation, we employed a cross-modality matching scheme to equalize the stimuli intensities of each modality according to the perception of an individual participant. Our results showed that the addition of both vibrotactile and joint-torque feedback improved object discrimination accuracy over no haptic feedback, but no difference was found between the two haptic feedback conditions themselves. These findings indicate that kinesthetic and vibrotactile feedback are equally beneficial methods of augmenting a traditional myoelectric prosthesis. Even though the task was force-based, cutaneous feedback performed the same as kinesthetic feedback.

## Ethics approval and consent to participate

All participants were consented according to a protocol approved by the Johns Hopkins School of Medicine Institutional Review Board (Study# IRB00147458), and were compensated at a rate of $10 per hour.

## Consent for publication

The individual featured in the images in this manuscript is the first author, who consents to this publication.

## Availability of data and materials

The datasets used and/or analysed during the current study are available from the corresponding author on reasonable request.

## Author’s contributions

NT and CM designed the experimental protocol. NT and GU ran the study. NT analyzed and interpreted the data. NT, GU, and JDB prepared the manuscript. All authors have reviewed and agreed to the final manuscript submission.

## Acknowledgements

The authors would like to thank Christopher Scherz and Jacob Carducci for their help in designing, building and maintaining the actuation system. We would also like to thank Dankmeyer, Inc. for providing the prosthesis components and Leah Jager from the Johns Hopkins Institute for Clinical and Translational Research for providing statistical analysis consulting services.

